# Prenatal Dexamethasone Programs Autonomic Dysregulation in Female Rats

**DOI:** 10.1101/2024.08.05.606452

**Authors:** Lakshmi Madhavpeddi, Monique Martinez, Jared Alvarez, Arpan Sharma, Chengcheng Hu, Stuart A Tobet, Taben M Hale

## Abstract

Autonomic dysfunction is associated with cardiovascular and neurological disease, including hypertension, heart failure, anxiety, and stress-related disorders. Prior studies demonstrated that late gestation exposure to dexamethasone (DEX) resulted in female-biased increases in stress-responsive mean arterial pressure (MAP) and heart rate (HR), suggesting a role for glucocorticoid-mediated programming of autonomic dysfunction. The present study investigated the influence of sympathetic (SYM) or parasympathetic (PS) blockade on cardiovascular function in male and female rat offspring of mothers injected with DEX *in utero* (gestation days [GD]18-21). At 11-12-weeks of age, MAP, HR, and heart rate variability (HRV) were evaluated at baseline and in response to SYM antagonists (α_1_-adrenoceptor + β_1_-adrenoceptor), a PS (muscarinic) antagonist, or saline (SAL). To assess stress-responsive function, rats were exposed to acute restraint. Tyrosine hydroxylase was measured in adrenals and left ventricle, and gene expression for the β_1_ adrenergic receptor was measured in left ventricle. Maternal DEX injection reduced basal HRV in male and female offspring. SYM blockade attenuated increases in stress-responsive HR and MAP. PS blockade elevated stress-responsive HR and MAP to a greater extent in Vehicle females. SYM and PS blockade produced equivalent effects on HR and MAP responses in male offspring, regardless of maternal treatment. Based on these findings, we suggest that maternal DEX injection disrupted autonomic regulation of cardiovascular function in females, resulting in a shift toward greater SYM input and less input from PS. Future studies will investigate whether changes in autonomic function are mediated by changes in central autonomic circuitry.

**New and Noteworthy:** These studies use pharmacological antagonists to characterize the nature of the autonomic dysregulation induced in female offspring exposed to the synthetic glucocorticoid, dexamethasone, *in utero*. The female offspring of dams injected with dexamethasone in late gestation show a reduction in vulnerability to parasympathetic blockade and an increase in responses to acute restraint stress even in the presence of sympathetic blockade. This suggests that late gestation dexamethasone disrupts the normal development of the autonomic function in females leading to a shift in the sympathovagal balance.

## Introduction

Autonomic dysregulation is a known independent risk factor for hypertension, arrhythmia, acute myocardial infarction, heart failure and stroke (1-5). Studies focused on the developmental origins of disease have shown prenatal perturbations can increase risk for development of cardiovascular disease and other chronic adult-onset diseases, and autonomic dysfunction may underlie these effects. Prenatal conditions leading to elevated glucocorticoids *in utero* have been shown to alter cardiovascular, neuroendocrine, and behavioral responses at baseline and in response to stress in rodent studies (6-11). Studies in our laboratory have demonstrated that adult rat offspring from dams injected with the synthetic glucocorticoid dexamethasone (DEX) during the last four days of gestation exhibit female-selective elevations in core body temperature, anxiety-like behaviors, as well as stress-responsive blood pressure and heart rate, suggestive of autonomic dysregulation (8-11). The synthetic glucocorticoids, dexamethasone or betamethasone have been routinely administered to mothers at risk for preterm labor to promote fetal lung development, and to COVID-positive mothers with need for supplemental oxygen (12-16). The long-term consequences of this exposure on development remain an important area for investigation.

Cardiac autonomic dysregulation is an imbalance of sympathetic and/or parasympathetic regulation of basal and stress-evoked cardiovascular function and is commonly associated with an overactive sympathetic nervous system and elevated circulating catecholamines (17-20). Heart rate variability (HRV) measures the interval between successive heart beats and is an established method for non-invasive measurement of autonomic regulation and sympathovagal balance clinically and preclinically. Low heart rate variability is associated with increased morbidity and mortality related to hypertension, arrhythmia, coronary artery disease, diabetes as well as depression and anxiety (5, 21, 22).

Females have been shown to have higher parasympathetic drive and lower sympathetic input, compared to males, which may contribute to lower incidence and later onset of cardiovascular disease in premenopausal women (3-25). In rats, maternal administration of the synthetic glucocorticoid, DEX, results in an altered pressor and tachycardic responses to acute stress in female adult offspring, suggesting altered cardiac autonomic regulation (11). We hypothesize that elevations in sympathetic activation or withdrawal of vagal input contribute to the altered physiological profile programmed by maternal DEX injection. The objective of this study is to determine the extent to which maternal DEX injection changes the sympathetic (SYM) or parasympathetic (PS) activity in the adult offspring by measuring arterial pressure and heart rate following administration of (1) α- and β-adrenergic antagonists to block the effects of SYM activation and (2) muscarinic antagonism to block the effects of PS activation. These effects were evaluated at rest and in response to an acute stressor. Heart rate variability was measured at rest to evaluate the degree to which maternal exposure to DEX altered autonomic function.

## Methods

### Animals

Sprague Dawley rats (Charles River Laboratories, Wilmington, MA) were bred in house, and single housed beginning at gestation day (GD) nine with ad libitum food and water in a temperature (70-75 °F) and humidity (30-50%)-controlled environment on a standard 12:12 light/dark cycle (lights on at 6am and off at 6pm). Dams were handled daily for 5 minutes on GD 14-17 to acclimate rats and minimize stress associated with experimental manipulations. Dams were randomly assigned to receive either DEX (n= 12) or vehicle (n=12). As outlined in Figure 1, DEX (0.4 mg/kg/day, s.c.) or vehicle (hydroxypropyl beta cyclodextrin; Veh) was administered daily on GD 18-21, as previously described (11, 26). Litters were reduced to 10 (approximately 5 males and 5 females per litter) to minimize postnatal nutritional variability. Pups were weaned at 21 days and group housed by sex (27). Rats were housed at a minimum of 2 per cage until they reached a body weight that impeded pair housing. Single housed rats were provided IACUC approved enrichment. All assessments were performed in male and female adult offspring (11-12 weeks old); with a maximum of one male and one female randomly chosen from each litter. All procedures were approved by the University of Arizona Institutional Animal Care and Use Committees and cared for in accordance with recommendations in the National Institutes for Health Guide for the Care and Use of Laboratory Animals, 8th Edition (NIH Pub. No. 85-23, Revised 2011).

**Figure 1.**
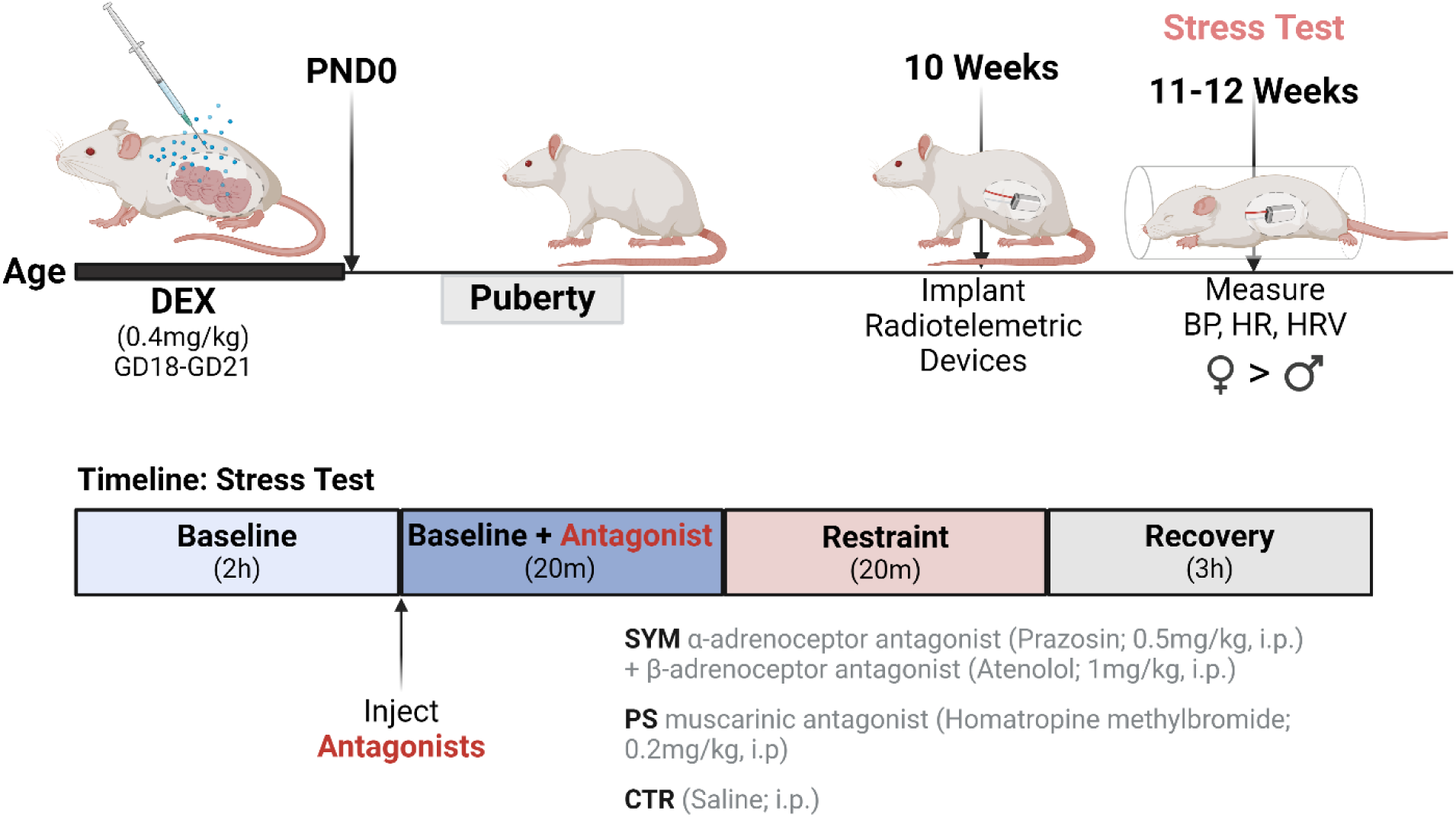
Timeline for experimental protocol. Created with BioRender.com and published with permission.

### Radiotelemeter Implantation

Radiotelemetric devices (Model – HD-S10; Data Sciences International (DSI), St. Paul, MN) were implanted into the abdominal aorta of adult male and female animals at approximately 10 weeks of age, as previously described (2, 24, 25). Buprenorphine SR (1mg/kg, s.c., ZooPharm, Windsor, CO) was administered 30-45 minutes prior to the start of surgery for general analgesia, and Lidocaine (Med-Vet International, Mettawa, IL) was applied to the skin suture line post-surgery for topical analgesia. Clavamox (Zoetis, Parsippany, NJ) was administered daily for four days to prevent infection. Following radiotelemeter implantation, animals were single housed in a dedicated radiotelemetry housing room on a 14:10 light/dark cycle (lights on at 6AM and off at 8PM) and allowed to recover for 1 week prior to experimental manipulations. Following recovery, estrous cycle stages were determined by daily vaginal lavage and cytological analysis.

### Telemetry, Measurement of Cardiovascular Parameters

Cardiovascular parameters, including mean arterial pressure (MAP), heart rate (HR) and heart rate variability (HRV) were collected throughout the experiment at a sampling rate of 500Hz using Ponemah software (DSI, St. Paul, MN; Version 10.0). HRV was calculated with Ponemah software, using the interbeat interval acquired from blood pressure waveforms. Frequency domain analysis of HRV was evaluated, and data integrated into low frequency (LF: 0.20-0.75Hz) and high frequency (HF: 0.75-2.5Hz) bands (28, 29). HF has been shown to correspond parasympathetic control, whereas LF has been described to contain input from both parasympathetic (PS) and sympathetic (SYM) (28, 30, 31). Time domain analysis of HRV was analyzed for root mean square of successive differences (RMSSD), a measure of vagal input, and standard deviation of N-N interval (SDNN) for successive heart beats (30).

### Impact of Pharmacological Antagonists on restraint stress

On the day of testing, the frequency and segment duration of data collection was set to 1 minute every 5 minutes. Basal MAP, HR, and HRV were measured for 20 minutes, prior to experimental testing. To evaluate the influence of SYM vs. PS inputs on cardiovascular responses to restraint stress, rats were administered α-adrenoceptor (prazosin; 0.5mg/kg, *i.p.*) and β-adrenoceptor (atenolol; 1mg/kg, *i.p.*) antagonists to attenuate SYM activity, a muscarinic receptor antagonist (homatropine methylbromide; 0.2mg/kg, *i.p.*) to block peripheral PS activity, or saline control (SAL) (32). The SAL group was used as a control for comparison to assess the impact of prenatal DEX exposure on SYM responses and to assess the impact of prenatal DEX exposure on PS responses. Using the same control group to address these distinct assessments allowed for the reduction in the total number of rats required for this assessment, in line with the 3 Rs principle in animal research. Rats were randomly assigned to a given treatment group. Twenty minutes following injection of pharmacological antagonists or saline, rats were placed in a plexiglass restraint tube in their home cage for 20 minutes. This timeline was based on the return of MAP to baseline following saline injection in a small pilot study. Females were tested on diestrus. Baseline MAP, HR, and HRV were determined based on the average of 5 data points preceding the antagonist or saline administration. The delta change was calculated from baseline following injection of antagonists and during restraint. Three hours after stress testing, rats were euthanized by CO_2_ inhalation followed by decapitation. Left ventricle and adrenal glands were extracted from a separate cohort of age-matched rats, flash frozen, and stored on at - 80°C until use.

### Real-time quantitative PCR

RNA was isolated from LV tissue using Trizol (Ambion, 15596018) as per the manufacturer’s instructions. cDNA was synthesized using reverse transcriptase (Invitrogen, 28025-013) with oligo-dT using a BioRad C1000 Thermal Cycler. Real-time PCR was performed with an Applied Biosystems QuantStudio 6 PCR system using SYBR green (Quanta, 95074-012). Suitably designed primers were ordered from IDT (Coralville, IA) for adrenergic receptor β_1_ (*Adrb1*) (*Adbr1* Fwd: CGCTGCCCTTTCGCTACCAG; Rev: CCGCCACCAGTGCTGAGGAT) and the housekeeping gene Glyceraldehyde 3-phosphate dehydrogenase (*Gapdh* Fwd: AGTGCCAGCCTCGTCTCATA; Rev: GGTAACCAGGCGTCCGATAC). The PCR conditions were as follows: 50°C for 2 min, 95°C for 10 min, 40 cycles of 95°C for 15 sec, and 60°C for 1 min. Product specificity was verified with a melt curve and by running real-time PCR products on a 1% agarose gel containing ethidium bromide.

### Immunoblotting

Protein quantification was performed using a standard western blotting procedure for tyrosine hydroxylase (TH 1:800, Abcam ab112). Samples from males and females were run in separate gels. Thus, sex differences for these assessments were based on the degree to which maternal DEX injection produced a different effect in each sex. Protein levels were normalized to total protein within the lane (Revert™ 700 Total Protein Stain Kit, Licor, 926-11016). Data was normalized to average Vehicle within sex. A subset of LV and adrenals from vehicle male and female rats were run on a single gel to evaluate the degree to which there are basal sex differences in TH expression.

### Statistical Analysis

Impact of maternal DEX injection on maternal body weight was analyzed by an unpaired Student’s t-test (GraphPad Prism 10). Analysis of birth weight, baseline MAP, HR, and HRV, as well as protein and mRNA levels were assessed by Two Way ANOVA with maternal injection and sex as the variables. For HRV in the frequency domain, log transformation was performed prior to performing two-way ANOVA. Prior to statistical analysis of HRV, all blood pressure waveforms were evaluated in a blinded fashion using the Ponemah software. Analysis included marking of physiologically relevant waveforms that were missed by the software, and removal of non-physiological waveforms (e.g., noise and skipped beats) and data breaks for which no waveforms were detected. Any segments (i.e. 1 min time period) with a data break or removal of waveforms that exceed a 500msec threshold was excluded by the software for frequency domain analysis. For HRV in the frequency domain, differences in variances across groups necessitated log transformation prior to performing two-way ANOVA (GraphPad Software). For HRV in the time domain, analysis was performed on raw values. Protein and mRNA expression data were normalized to the average of the vehicle group within sex to evaluate the degree to which prior DEX impacted expression in a sex-specific manner.

To assess the impact of pharmacological antagonist and restraint stress on MAP and HR, a linear mixed effects model was fitted for changes in each marker (MAP or HR) from baseline (average of five measurements) to the five time points in each of the two time periods (phases) under study (baseline with antagonist and stress), adjusting for baseline level, time point (five-level factor, with the first time point as reference, or a linear time since start of the phase in minutes), a two-level factor of in-utero intervention (dexamethasone, or DEX, versus vehicle only, or Veh), a two level factor of antagonist (PS vs. SAL for studies of PS effects, or SYM vs. SAL for studies of SYM effects; each antagonist was studied separately with rats on the other antagonist excluded), and interaction of in-utero intervention and the antagonist. When the interaction term was significant at the 0.05 level, effect of each antagonist (vs SAL) was reported separately for each group of in-utero intervention (DEX or Veh); if interaction was not significant the model was re-fitted with that term removed and overall DEX effect and antagonist effect were reported. We explored whether DEX or each antagonist modified the slope of change over time by using time since start of the specific phase as a continuous variable in the model and including an interaction term between DEX/antagonist and the continuous time. An animal-level random intercept was included to account for correlation of multiple measurements (five in each phase) on the same rat, and the within-animal correlation structure was assumed to be the autoregressive process with order 1. All statistical tests were two-sided with significance level 0.05. The statistical software environment R (version 4.3.3 is used for all analysis (33). The R package nlme was used to fit linear mixed effect models (34). Males and females were analyzed separately.

In all cases, a p < 0.05 was considered statistically significant and data are presented as mean ± standard error of the mean (SEM).

## Results

### Maternal Injection of DEX Promoted Growth Restriction

At the time of initiating maternal injections (GD18), there was no difference in maternal body weight between dams assigned to the two treatment groups (Veh: 350 ±10.5g, DEX: 343 ± 9.2g, p=0.62). However, at GD21 the body weight of DEX injected dams was significantly lower than those of vehicle injected dams (Veh: 388 ± 10.5g, DEX: 349 ± 9.8g, p=0.013). There were no differences in the sizes of the litters, prior to culling to 10 per litter (Veh: 14 ± 0.5, DEX 14 ± 0.7 pups, p = 0.9), nor was there an impact on the ratio of males to females in a given litter (Veh: 1 ± 0.2, DEX 1 ± 0.1 males relative to females, p=0.3). Maternal DEX injection resulted in a significant reduction in pup birth weight (Veh male: 6.6 ± 0.11g, Veh female: 5.9 ± 0.19g, DEX male: 5.8 ± 0.15g, DEX female: 5.4 ± 0.2g; main effect of maternal injection: p = 0.0009; main effect of sex: p = 0.0047; maternal injection x sex: p = 0.25).

### Impact of prenatal DEX on basal HRV, HR, and MAP

Maternal DEX injections altered baseline HRV both in female and male rat offspring in adulthood, as assessed by frequency and time domain analyses. Prenatal DEX exposure resulted in a lower log(LF) power (p = 0.0255 DEX vs. Veh; Fig. 2A), which is suggested to reflect both SYM and PS input, and log(HF) power (p 0.0173 DEX vs. Veh; Fig. 2B), an index of vagal tone. Similarly, prenatal exposure resulted in a lower SDNN (p = 0.0079 DEX vs. Veh; Fig. 2C) and RMSSD (p = 0.0038 DEX vs. Veh; Fig 2D). Prenatal DEX did not significantly alter baseline HR in female or male rats (Female Veh: 342 ± 5.9 bpm vs. DEX: 355 ± 8.2 bpm; Male Veh: 310 ± 6.8 bpm vs. DEX: 315 ± 7.4 bpm), but there was a main effect of sex with females having higher HR than males (Veh: p = 0.0111 Male vs. Female; DEX: p = 0.0010 Male vs. Female). Prenatal DEX did not significantly alter baseline MAP in female or male offspring (Female Veh: 108 ± 1.0 mmHg vs. DEX: 107 ± 1.8 mmHg, p >0.05. Male Veh: 105 ± 1.4 mmHg vs. DEX 108 ± 1.4 mmHg; p >0.05), nor was there an effect of sex.

**Figure 2.**
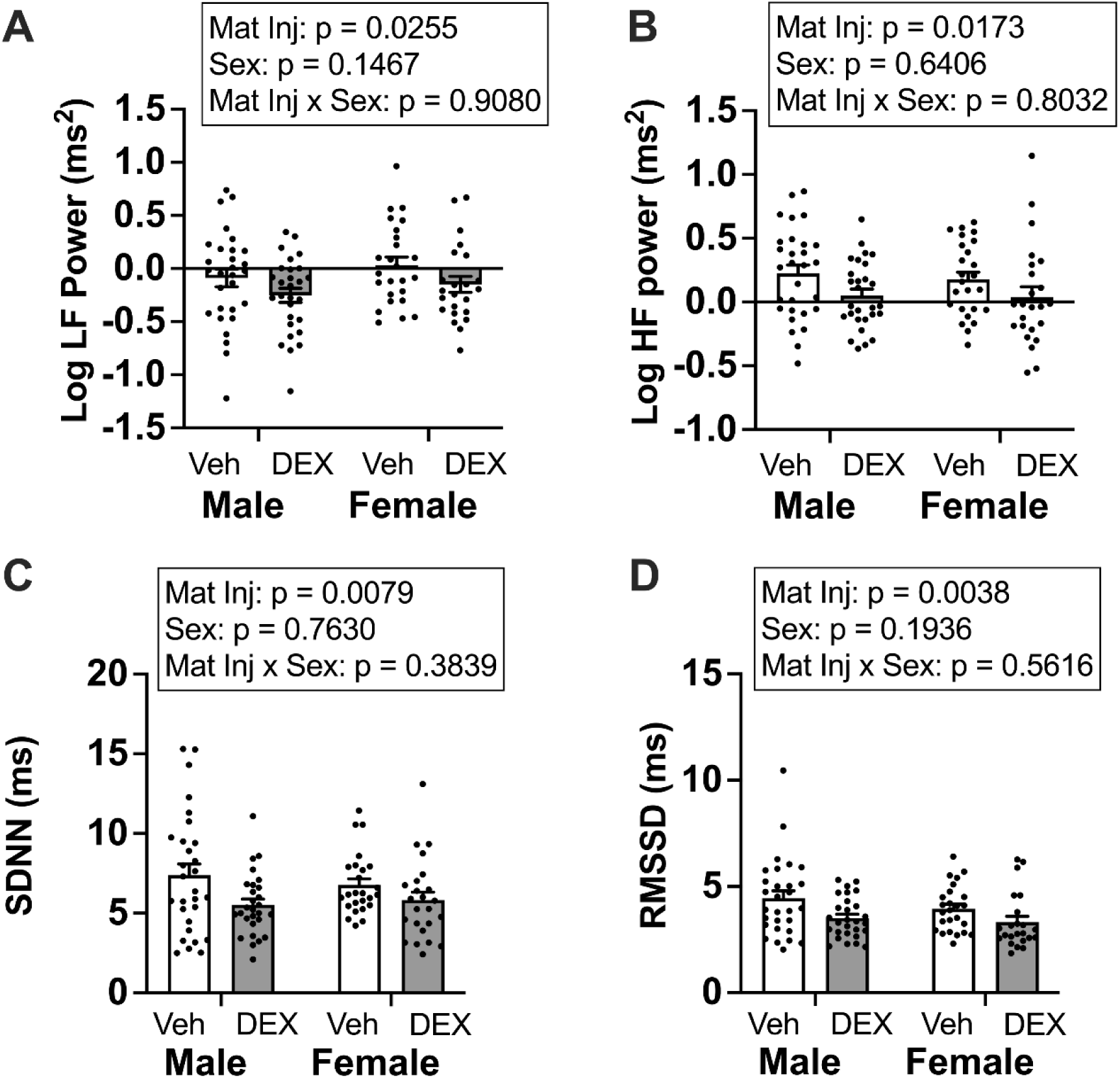
Assessments of log-transformed low frequency (LF) power (A), log-transformed high frequency (HF) power (B), standard deviation of heartbeat intervals (SDNN) (C) and the root mean square of successive differences (RMSSD) (D) in vehicle (Veh) and dexamethasone (DEX)-exposed male and female rats was evaluated by two-way ANOVA. Data presented as mean ± SEM, N=16-29 per group. Mat inj: maternal injection. Data panels were assembled in BioRender.com and published with permission.

### Impact of SYM blockade on basal and stress-evoked HR

HR responses to restraint stress in the presence of SYM blockade were assessed in female offspring (Fig. 3A, B, C). SYM blockade resulted in a lower HR response to *i.p.* injection that resulted in a significant main effect of the antagonist (p < 0.0001; Fig 3B). The rate of change in HR following bolus injection was significantly impacted by SYM antagonist (p=0.0048), but not by maternal injection. In response to restraint, the increase in HR was attenuated by SYM blockade regardless of maternal injection (main effect of antagonist: p < 0.0001; Fig 3C). HR responses to restraint were higher in DEX-exposed females even in the presence of SYM antagonists (main effect of maternal injection: p=0.0065; Fig3C). Although there was not a statistically significant interaction, there was an apparent greater effect of SYM blockade in DEX exposed females (Fig. 3C). The rate of HR recovery was significantly impacted by SYM (p=0.0001), but not by maternal injection.

**Figure 3.**
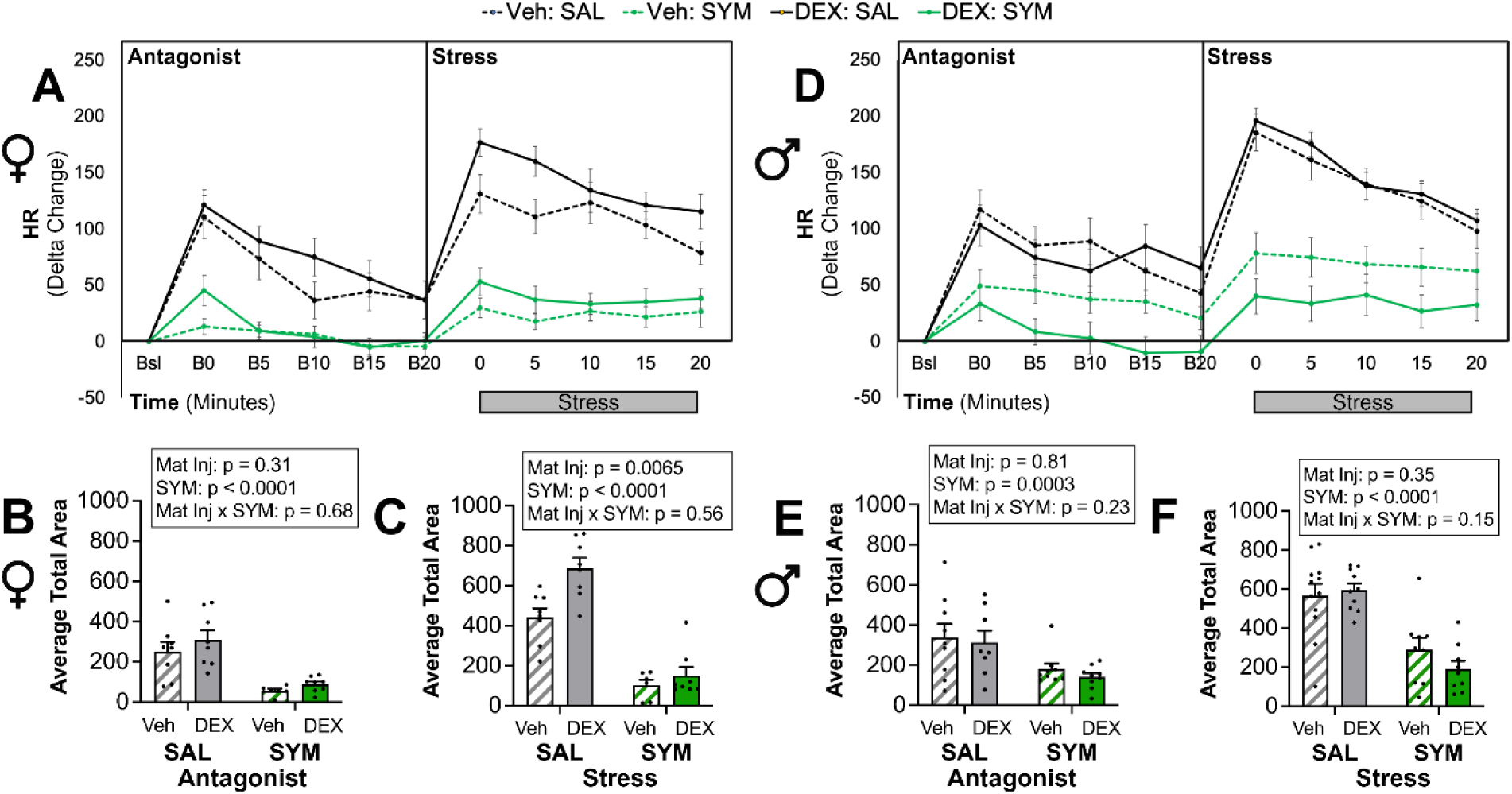
Impact of SYM Blockade on Heart Rate. Heart rate following injection of α-adrenoceptor (prazosin) and β-adrenoceptor (atenolol) antagonists to block peripheral sympathetic (SYM) activity and during restraint in female (A, B, C) male (D, E, F) offspring of dams exposed to vehicle (Veh) or dexamethasone (DEX). Line graphs (A and D) depict change from the average of five successive timepoints immediately prior to injection. Grey bar depicts the 20-minute restraint period. Bar graphs depict the average area under the curve for the baseline period following injection of antagonist or saline (SAL) in female (B) and male (E) offspring, and the period of restraint in female (C) and male (F) offspring. Statistics reported for each bar graph are based on mixed model analysis after controlling for baseline heart rates for each rat, as described in methods. Data presented as mean ± SEM, N=6-10 per group. Bsl: average baseline. B0, B5, B10, B15, B20: time in minutes following bolus (B) injection. Mat inj: maternal injection. Data panels were assembled in BioRender.com and published with permission.

HR responses to restraint stress in the presence of SYM blockade were assessed in male offspring (Fig. 3D, 3E, 3F). There was a main effect of SYM blockade to reduce the HR response to injection (main effect of SYM antagonism p = 0.0003) compared to SAL injected rats that was not influenced by prenatal DEX exposure. The rate of change in HR following bolus injection was not significantly impacted by SYM antagonist or by maternal injection. In response to restraint, the HR response was significantly attenuated by SYM blockade (main effect of antagonist: p < 0.0001; Fig. 3F). Although there was not a statistically significant main effect of maternal injection or an interaction, there was an apparent greater effect of SYM blockade in DEX exposed males. The rate of HR recovery was significantly impacted by SYM (p<0.0001), but not by maternal injection.

### Impact of PS blockade on basal and stress-evoked HR

Overall, PS antagonist treatment differentially impacted HR responses in VEH vs. DEX exposed females (Fig. 4A, B, C). At baseline, compared to SAL, PS blockade increased the HR response to the bolus injection (main effect of antagonist, p= 0.013; Fig. 4B). The rate of change in HR following bolus injection was significantly impacted by PS antagonist (p=0.0021), but not by maternal injection. During restraint, there was a significant maternal injection x PS antagonist interaction (p = 0.049) whereby PS blockade resulted in a marked increased HR response to restraint in Veh-exposed females and a decreased HR response to restraint in DEX-exposed females. Neither PS antagonism nor maternal injection influenced the rate of HR recovery to restraint.

**Figure 4.**
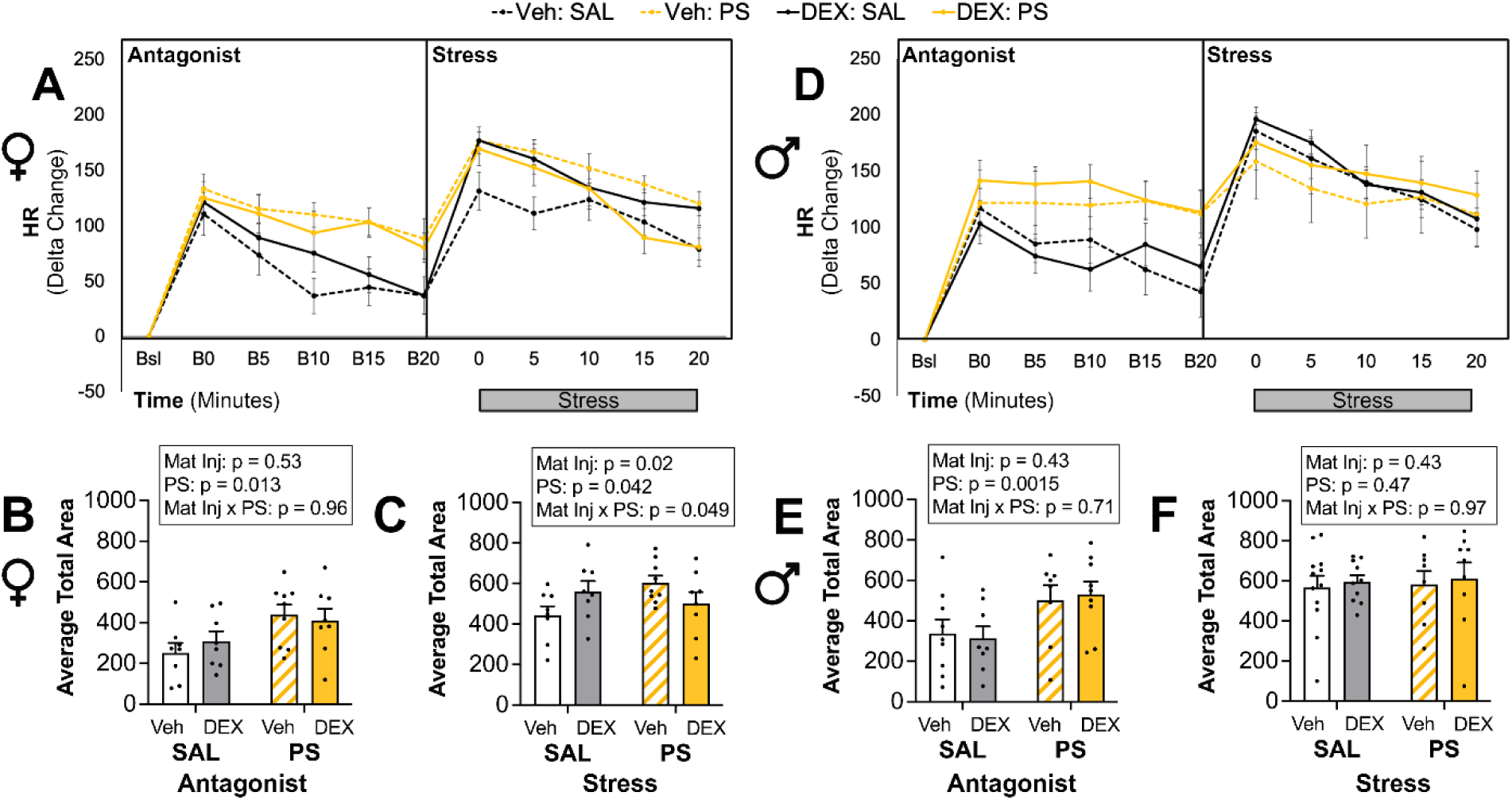
Impact of PS Blockade on Heart Rate. Heart rate following injection of a muscarinic receptor antagonist (homatropine methylbromide) to block peripheral parasympathetic (PS) activity and during restraint in female (A, B, C) male (D, E, F) offspring of dams exposed to vehicle (Veh) or dexamethasone (DEX). Line graphs (A and D) depict change from the average of five successive timepoints immediately prior to injection Grey bar depicts the 20-minute restraint period. Bar graphs depict the average area under the curve for the baseline period following injection of antagonist or saline (SAL) in female (B) and male (E) offspring, and the period of restraint in female (C) and male (F) offspring. Statistics reported for each bar graph are based on mixed model analysis after controlling for baseline heart rates for each rat, as described in methods. Data presented as mean ± SEM, N=6-10 per group. Bsl: average baseline. B0, B5, B10, B15, B20: time in minutes following bolus (B) injection. Mat inj: maternal injection. Data panels were assembled in BioRender.com and published with permission.

HR responses to restraint stress in the presence of PS blockade were assessed in male offspring (Fig. 4 D, E, F). There was a main effect of PS blockade (Fig. 4D) to increase the HR response to injection (main effect of PS antagonism p = 0.0015; Fig. 4E), with no influence of prenatal DEX exposure. The rate of HR change following bolus injection was significantly impacted by PS antagonist (p=0.015), but not by maternal injection. In response to restraint, PS blockade did not influence the HR response to restraint compared to SAL, regardless of prenatal DEX exposure (Fig. 4F). The rate of HR recovery to restraint stress was significantly influenced by PS blockade (p=0.0002), but not maternal injection.

### Impact of SYM blockade on basal and stress-evoked MAP

MAP responses to restraint stress in the presence of SYM blockade were assessed in female offspring (Fig. 5A, B, C). In females, *i.p.* injection with saline resulted in an increase in MAP (Fig. 5). Compared to SAL, SYM blockade resulted in a lower MAP response to bolus injection (main effect of antagonist, p < 0.0001; Fig 5B). The rate of change in MAP following bolus was significantly impacted by SYM antagonist (p=0.0005), without influence of maternal injection. In response to restraint, the increase in MAP was attenuated by SYM antagonism (main effect of antagonist: p <0.0001; Fig. 5C). Neither antagonist nor maternal injection influenced the rate of MAP recovery during restraint.

**Figure 5.**
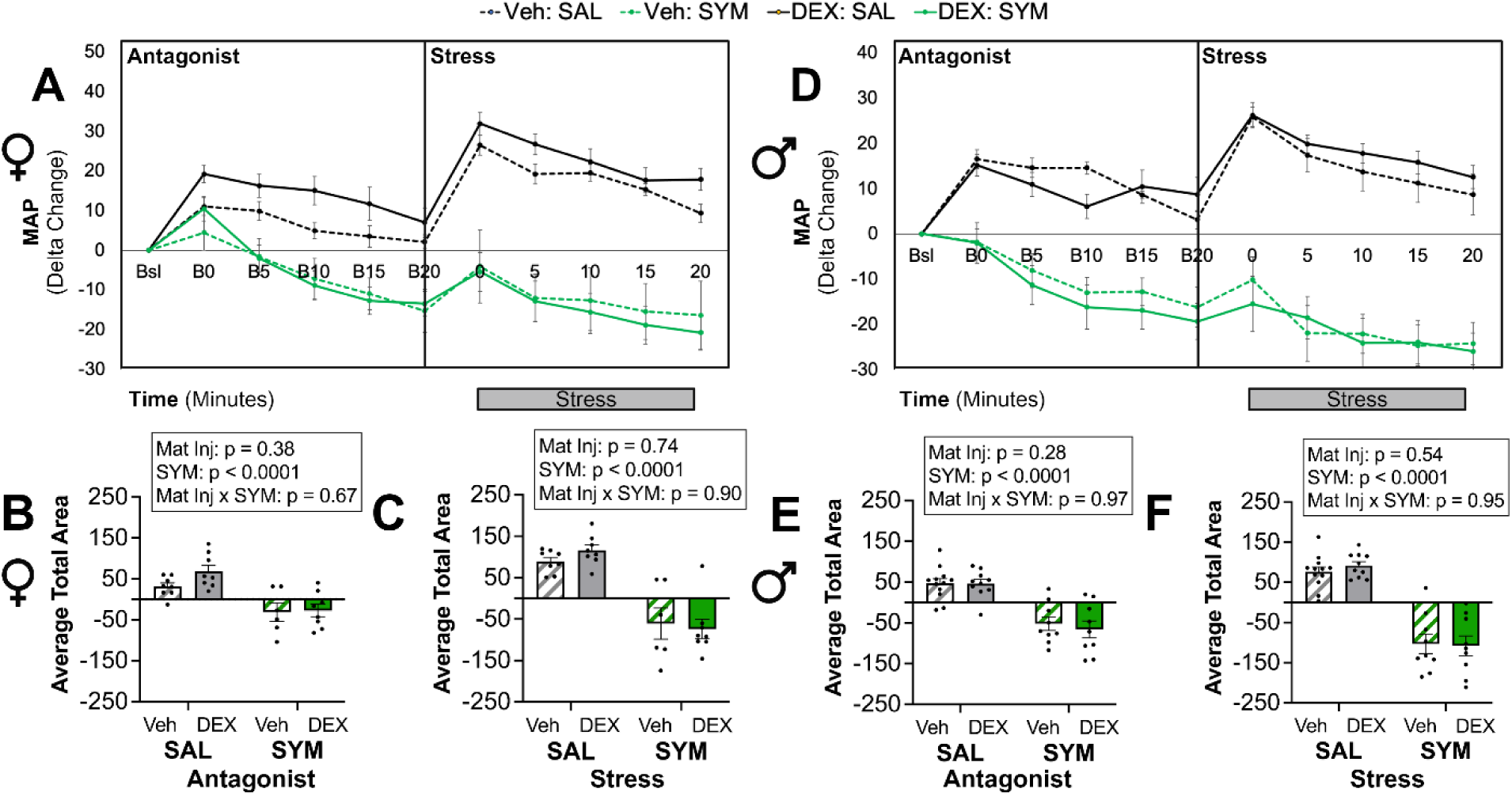
Impact of SYM Blockade on Blood Pressure. Blood pressure following injection of α-adrenoceptor (prazosin) and β-adrenoceptor (atenolol) antagonists to block peripheral sympathetic (SYM) activity and during restraint in female (A, B, C) male (D, E, F) offspring of dams exposed to vehicle (Veh) or dexamethasone (DEX). Line graphs (A and D) depict delta change from the average of five successive timepoints immediately prior to injection. Grey bar depicts the 20-minute restraint period. Bar graphs depict the average area under the curve for the baseline period following injection of antagonist or saline (SAL) in female (B) and male (D) offspring, and the period of restraint in female (C) and male (F) offspring. Statistics reported for each bar graph are based on mixed model analysis after controlling for baseline mean arterial pressures for each rat, as described in methods. Data presented as mean ± SEM, N = 6-10 per group. Bsl: average baseline. B0, B5, B10, B15, B20: time in minutes following bolus (B) injection. Mat inj: maternal injection. Data panels were assembled in BioRender.com and published with permission.

MAP responses to restraint stress in the presence of SYM blockade were assessed in male offspring (Fig. 5D, E, F). In response to saline injection there was an increase in MAP that was significantly attenuated by SYM antagonists (main effect of antagonist, p < 0.0001; Fig. 5E). The rate of change in MAP following bolus injection was significantly impacted by SYM antagonist (p=0.0055), but not by maternal injection. In response to restraint, the MAP response was significantly attenuated by SYM blockade (main effect of SYM antagonism: p < 0.0001; Fig. 5F), with no impact of maternal injection of DEX. The rate of MAP recovery during restraint was not significantly impacted by SYM antagonists or by maternal injection.

### Impact of PS blockade on basal and stress-evoked MAP

MAP responses to restraint stress in the presence of PS blockade were assessed in female offspring (Fig. 6A, B, C). PS blockade resulted in a greater increase in MAP in response to bolus injection (main effect of antagonist p = 0.0001; Fig 6B) – an effect that appeared to be more pronounced in Veh-exposed rats. PS blockade significantly influenced the rate of change in MAP following bolus injection (p=0.0099), without influence of the maternal injection. MAP was not further changed during restraint, regardless of prenatal exposure, although there was a tendency toward an increase in Veh-exposed offspring and decrease in DEX-exposed offspring (Fig. 6C).

**Figure 6.**
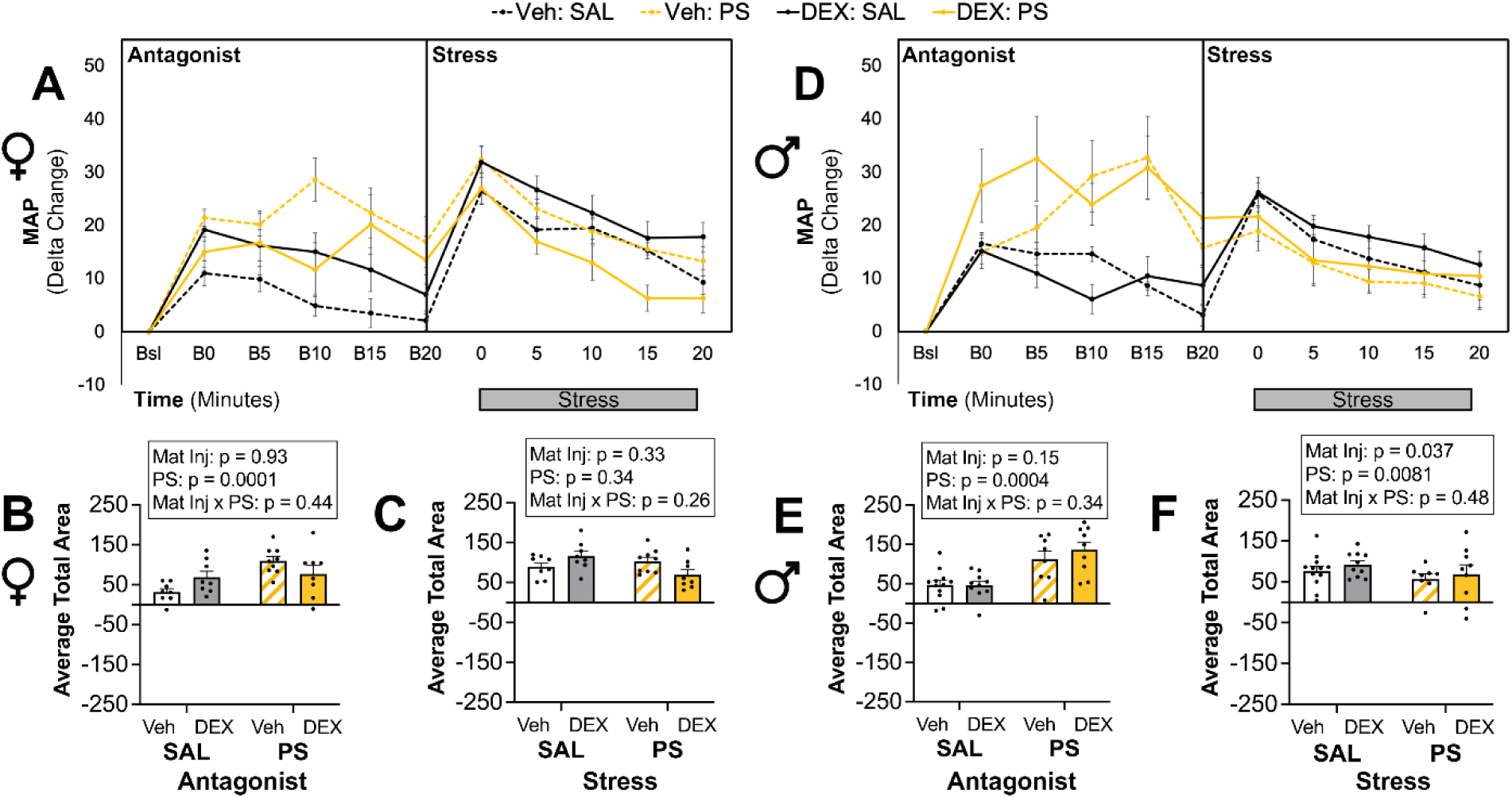
Impact of PS Blockade on Blood Pressure. Blood pressure following injection of a muscarinic receptor antagonist (homatropine methylbromide) to block peripheral parasympathetic (PS) activity and during restraint in female (A, B, C) male (D, E, F) offspring. Line graphs depict change from the average of five successive timepoints immediately prior to injection. Grey bar depicts the 20-minute restraint period. Bar graphs depict the average area under the curve for the baseline period following injection of antagonist or saline (SAL) in female (B) and male (E) offspring, and the period of restraint in female (C) and male (F) offspring. Statistics reported for each bar graph are based on mixed model analysis after controlling for baseline mean arterial pressures for each rat, as described in methods. Data presented as mean ± SEM, N = 6-10 per group. Bsl: average baseline. B0, B5, B10, B15, B20: time in minutes following bolus (B) injection. Mat inj: maternal injection. Data panels were assembled in BioRender.com and published with permission.

MAP responses to restraint stress in the presence of PS blockade were assessed in male offspring (Fig. 6D, E, F). In male rats, there was a main effect of PS blockade to increase the MAP response to injection (main effect of antagonist: p = 0.0004; Fig. 6E). There was no statistically significant influence of maternal injection nor PS antagonist on the rate of change in MAP in response to the bolus injection. In response to restraint there was a main effect of PS to attenuate the MAP response (p=0.0081) and an overall main effect of maternal injection (p=0.037) (Fig. 6F).

### Impact of maternal DEX injections on tyrosine hydroxylase expression and gene expression for the β1 adrenergic expression in adult offspring

Expression of tyrosine hydroxylase (TH), the rate limiting step in catecholamine synthesis, was not significantly different in the left ventricle of male or female offspring of DEX injected mothers versus vehicle (Fig. 7A). However, there was a modest, yet significant interaction between sex and prenatal exposure in tyrosine hydroxylase expression in the adrenal gland (p = 0.0250, Fig. 7B). Total protein blots are available in *Supplemental Figure 1 (*https://doi.org/10.6084/m9.figshare.26145364). Male and female vehicle rats had similar expression of TH in LV and adrenal gland, suggesting a lack of basal sex differences (*Supplemental Figure* 2: https://doi.org/10.6084/m9.figshare.26145364). *Adbr1* gene expression in left ventricle was not significantly influenced by prenatal exposure or by sex (Fig. 7C).

**Figure 7.**
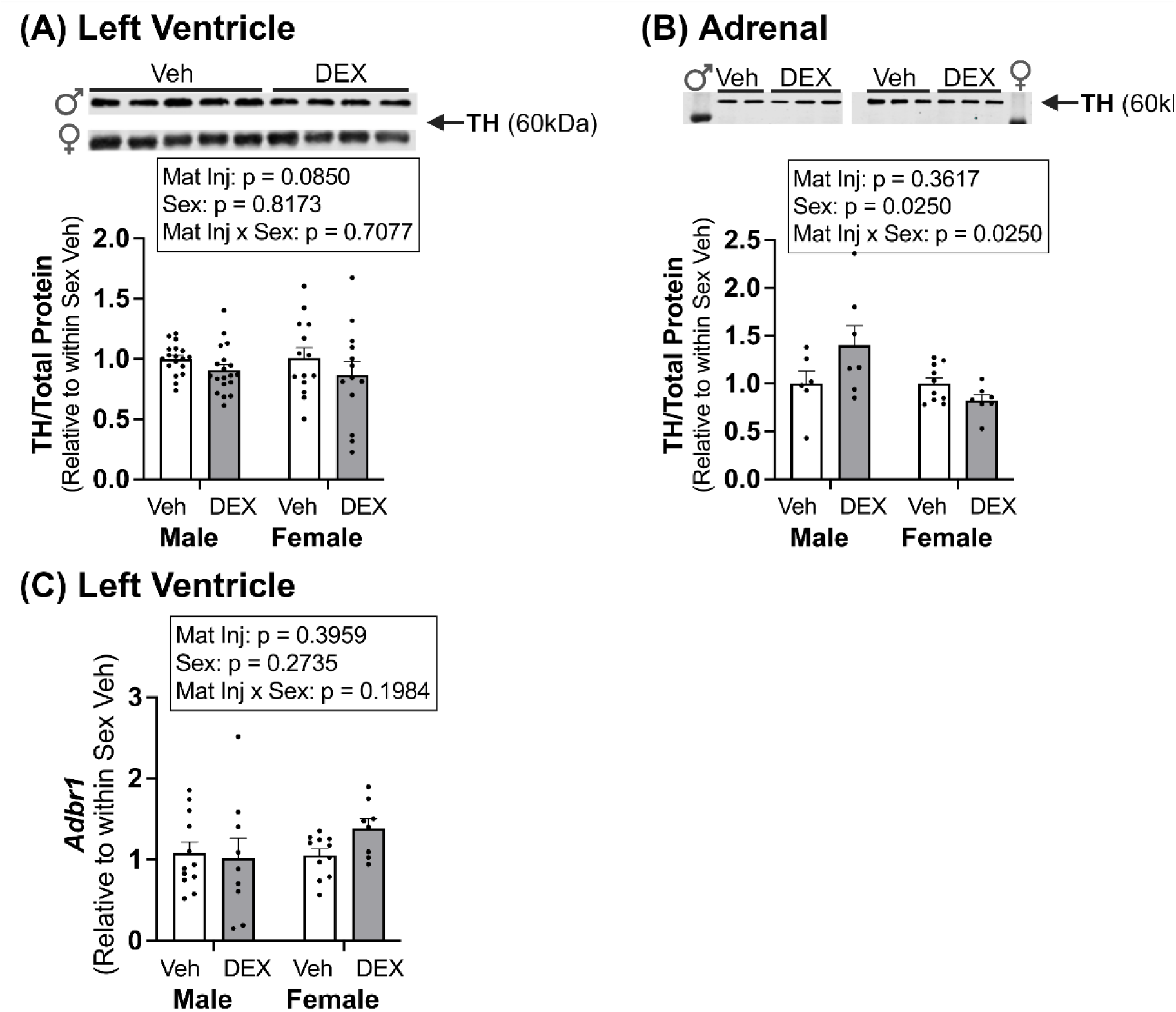
Protein and mRNA expression levels of enzymes or receptors involved in local catecholamine biosynthesis or activity in male and female offspring. Tyrosine hydroxylase (TH) protein expression levels in left ventricle N = 13-19 per group (A) and in adrenals N = 6-10 per group (B). Adrenergic receptor β_1_ (*Adbr1*) mRNA expression levels in left ventricle N = 8-12 per group (C). Data presented as fold change from average vehicle (Veh) mean ± SEM. DEX: dexamethasone. Mat inj: maternal injection. Data panels were assembled in BioRender.com and published with permission.

## Discussion

In the current study, injections of DEX to rat dams in the last 4 days of pregnancy resulted in altered autonomic function in their female offspring raised to young adulthood. This was characterized by changes in HRV, as well as MAP and HR responses to SYM or PS blockade in adult offspring. Specifically, (1) prenatal DEX reduced HRV, (2) SYM blockade attenuated the HR and MAP responses at baseline and in response to restraint stress, and this response tended to be greater in DEX-exposed animals, and (3) PS blockade increases the HR and MAP response at baseline and in response to restraint stress, and in females the impact is greater in Veh-exposed rats compared to DEX. These findings suggest that maternal DEX injections resulted in offspring with impaired autonomic development – particularly in females, resulting in a shift toward increased sympathetic and decreased parasympathetic tone.

As we have previously reported (11), maternal DEX injections did not result in altered basal MAP or HR in offspring but significantly elevated HR and MAP in response to restraint stress in females but not males of a similar age to the present study. These findings suggest that a secondary challenge in adulthood may be necessary to reveal glucocorticoid-induced programming effects on pressor and heart rate responses in females. This is consistent with primary data in a prenatal model of psychosocial stress and subsequent review of the literature across animal models of prenatal stress by Mastorci *et. al.* demonstrating that the impact of prenatal stress on cardiac autonomic function increases offspring susceptibility to a future stressor in a sex-specific manner and is supported by a metanalysis by Kajantie *et. al.* suggesting sex differences are not clear at baseline but revealed in response to psychosocial stress (35, 36).

Although HR and MAP are not different at baseline, maternal DEX injections altered basal HRV in male and female offspring. DEX exposure *in utero* reduced HRV indexes of vagal input, suggesting fetal exposure promotes autonomic dysregulation in adulthood, and this dysfunction may underlie the stress-responses changes in cardiovascular function we observed in female offspring.

The present studies used pharmacological antagonists to evaluate the degree to which maternal DEX injections resulted in altered responses in adult offspring. Specifically, HR and MAP responses to SYM and PS blockade were evaluated at rest and in response to restraint stress. We hypothesized that sympathetic hyperactivity and blunted vagal activity contribute to exaggerated stress responses in DEX-exposed female offspring.

SYM blockade (α- and β_1_-adrenergic receptor antagonists) significantly attenuated the HR and MAP responses to injection and to acute restraint in male and female offspring. Although there was not a significant interaction, we showed that SYM antagonism (adrenergic receptor antagonists) appeared to attenuate the increase in HR in response to restraint stress to a greater degree in DEX-exposed animals compared to Veh, suggesting that there is potentially a greater influence of SYM during stress on female offspring from DEX-injected mothers.

PS blockade (muscarinic receptor antagonist) significantly elevated the HR and MAP response to i.p. injection in male and female offspring. Prenatal DEX did not alter HR or MAP responses to PS blockade in male offspring. In females, we showed that PS blockade had a greater effect to increase HR and MAP responses to restraint stress in Veh-exposed females compared to DEX-exposed. These findings suggest that in healthy females (offspring of Veh-injected mothers) the parasympathetic nervous system serves to attenuate the cardiovascular responses to stress. This response was disrupted in offspring exposed to DEX, *in utero.* Clinical studies show that women at rest exhibit higher HF power and lower LF power than men, suggestive of greater vagal regulation (23, 24, 37-41). Carnevali *et. al.* showed a similar finding in rodents, and further demonstrated that sex differences in HR and HRV are comparable between rodent and human models (28). Sex differences in cardiac autonomic control and heart rate in response to acute stress in humans are less clear.

Systemic reviews of clinical studies evaluating sex differences in response to acute psychosocial stress by Kajantie *et. al.* and Hamidovic *et. al.* found that in females, sympathoadrenal responsiveness, characterized by catecholamine levels, and HRV is reduced in females compared to males (36, 42) suggesting altered autonomic regulation in females. However, sex differences are not consistently described in clinical literature. Kajantie *et. al* suggests that differences may be due to sex-biased responses to the type of stressor, or in females, estradiol status and age (36). Prenatal programming effects on sex differences in cardiac autonomic responses to stress later in life remains an area for important future investigation.

Compared to Veh, DEX females exhibited decreased vulnerability to PS blockade, suggesting that prenatal exposure disrupts normal autonomic function. Taken together, female-specific alterations in HR and MAP in response to restraint following pharmacological intervention suggest that maternal injection of DEX shifts sympathovagal balance towards a sympathetic dominance, with less input from the parasympathetic nervous system in the offspring examined after puberty. This shift in autonomic control may be a risk factor for future cardiovascular disease.

The changes in cardiovascular responses in DEX-exposed females may be due changes in catecholamine bioavailability and increased activation of adrenergic receptors in the heart and the vasculature. Nguyen *et. al.* and Lamothe *et. al.* demonstrated elevated levels of circulating epinephrine and increased expression levels of adrenal enzymes involved in catecholamine synthesis in offspring of dams injected with DEX (43-45). Studies by Piquer *et. al.* and Jevjdovic *et. al.* showed decreases in β_1_ adrenergic receptor expression levels in the heart in response to prenatal chronic cold stress or chronic unpredictable mild stress (46, 47). We evaluated tyrosine hydroxylase (LV and adrenals) and β_1_ adrenergic receptor (LV) expression levels to determine whether they were altered in offspring of dams injected with DEX. β_1_ adrenergic receptor mRNA expression did not differ in the left ventricle. Moreover, protein expression levels of tyrosine hydroxylase (TH), the rate limiting enzyme in the production of norepinephrine, was not different in the left ventricle, though there was a trend towards an overall reduction in offspring of DEX-injected mothers (main effect of DEX: p = 0.0850). In the adrenals, prenatal DEX resulted in a modest elevation in TH protein expression levels in male rats, but slightly reduced expression levels in female rats. Overall, the changes in local regulation of catecholamine synthesis do not explain the observed functional responses. Changes in circulating levels of catecholamines, expression of additional synthesis and degradation enzymes for norepinephrine/acetylcholine, nerve density, receptor density and localization, as well as electrophysiological responses will be future areas of focus as potential peripheral changes induced by prenatal stressors.

The findings of the current study in males differ from prior studies (44, 45) showing increases in circulating levels of norepinephrine in DEX-exposed offspring, as well as male-specific increases in adrenal TH expression in DEX-exposed offspring, compared to Veh, as well as increases in downstream enzymes dopamine β-hydroxylase and phenylethanolamine n-methyl transferase (PNMT). Similarly, Nguyen and Khurana *et. al. s*howed increases in expression of adrenal PNMT and transcriptional activators Egr-1 and glucocorticoid receptor (GR) in males from DEX-injected mothers, and more recently demonstrated increases in adrenal TH, dopamine β-hydroxylase (DBH) and PNMT expression that are dose and sex dependent (43, 48). They found that increases in enzyme expression were observed only following a higher dose of DEX in the mothers of female offspring and suggest that females may be more resilient to prenatal DEX than males. These differences from our findings may be related to differences in the timing of DEX exposure. Specifically, in these abovementioned studies, DEX was administered from days 15 through 21 – a period that includes the prenatal testosterone surge. This earlier exposure to DEX has been shown to have a greater impact on male vs. female offspring. The timing of the DEX administration in our studies was chosen to more closely align with the clinical use of this drug (12). Moreover, it may be that the increased vulnerability to SYM withdrawal and reduced vulnerability to PS blockade we observe in female offspring of DEX-injected mothers in the present study is not due to autonomic control in the periphery but rather is regulated by central mechanisms. Future studies will evaluate the degree to which changes in autonomic control centers in the brain underlie DEX-dependent changes in cardiovascular function.

There are some limitations in the current study that should be noted. For example, longer continuous recordings may reduce the variability in HRV assessments observed in the present study. In addition, the timing of restraint post-antagonist injection was based on a pilot study in which we evaluated return of MAP to baseline for vehicle-exposed offspring. However, in some animals, HR remained elevated even 20 minutes post-saline injection. It is possible that this influenced the subsequent response to restraint. The pharmacological antagonists used in the present study were chosen to limit penetration of the blood brain barrier. This allowed for localization of the responses to the end organs that resulted from acetylcholine and norepinephrine release from nerve terminals and epinephrine release from adrenals. However, it is possible that there are central influences that are not accounted for in this design. In fact, our primary conclusion is that this is likely the case and will be the focus of future study.

In summary, the present findings demonstrate that maternal injection of DEX alters autonomic regulation of cardiovascular function in offspring. SYM or PS antagonists reveal a female-biased effect to shift sympathovagal balance from parasympathetic to sympathetic predominance. Autonomic dysregulation may underlie the female-specific exacerbations in HR and MAP in response to stress. Given that prenatal stress and glucocorticoid exposure during gestation are associated with cardiovascular disease in adults, identification of risk factors, such as altered autonomic function, is important for earlier intervention and prevention of future disease. Future studies will evaluate the role of central autonomic circuitry in mediating this sex-specific autonomic dysregulation.

## Supplemental Material

Supplemental Figures: https://doi.org/10.6084/m9.figshare.26145364

## Acknowledgements

This work was supported by the National Institutes of Health NIMH U54 MH118919 (TMH and ST), by NIMH U54 MH118919 S3 (MM), and by 5R25NS107188-05 (JA). The statistical analysis by CH is supported by the College of Medicine-Phoenix Biostatistics and Study Design Service, University of Arizona, College of Medicine – Phoenix.

## References

1 Tsuji H, Larson MG, Venditti FJ, Jr., Manders ES, Evans JC, Feldman CL, and Levy D. Impact of reduced heart rate variability on risk for cardiac events. The Framingham Heart Study. Circulation 94: 2850–2855, 1996.

2 Shen CJ, Tsou YA, Chen HL, Huang HJ, Wu SC, Cheng WT, Chen CY, and Chen CM. Osteoponin promoter controlled by DNA methylation: aberrant methylation in cloned porcine genome. Biomed Res Int 2014: 327538, 2014.

3 Bairey Merz CN, Elboudwarej O, and Mehta P. The autonomic nervous system and cardiovascular health and disease: a complex balancing act. JACC Heart Fail 3: 383–385, 2015.

4 La Rovere MT, and Christensen JH. The autonomic nervous system and cardiovascular disease: role of n-3 PUFAs. Vascul Pharmacol 71: 1–10, 2015.

5 Thayer JF, Yamamoto SS, and Brosschot JF. The relationship of autonomic imbalance, heart rate variability and cardiovascular disease risk factors. Int J Cardiol 141: 122–131, 2010.

6 Igosheva N, Klimova O, Anishchenko T, and Glover V. Prenatal stress alters cardiovascular responses in adult rats. J Physiol 557: 273–285, 2004.

7 O’Regan D, Kenyon CJ, Seckl JR, and Holmes MC. Glucocorticoid exposure in late gestation in the rat permanently programs gender-specific differences in adult cardiovascular and metabolic physiology. Am J Physiol Endocrinol Metab 287: E863–870, 2004.

8 Carbone DL, Zuloaga DG, Lacagnina AF, McGivern RF, and Handa RJ. Exposure to dexamethasone during late gestation causes female-specific decreases in core body temperature and prepro-thyrotropin-releasing hormone expression in the paraventricular nucleus of the hypothalamus in rats. Physiol Behav 108: 6–12, 2012.

9 Hiroi R, Carbone DL, Zuloaga DG, Bimonte-Nelson HA, and Handa RJ. Sex-dependent programming effects of prenatal glucocorticoid treatment on the developing serotonin system and stress-related behaviors in adulthood. Neuroscience 320: 43–56, 2016.

10 Eberle C, Fasig T, Brüseke F, and Stichling S. Impact of maternal prenatal stress by glucocorticoids on metabolic and cardiovascular outcomes in their offspring: A systematic scoping review. PLoS One 16: e0245386, 2021.

11 Madhavpeddi L, Hammond B, Carbone DL, Kang P, Handa RJ, and Hale TM. Impact of angiotensin II receptor antagonism on the sex-selective dysregulation of cardiovascular function induced by in utero dexamethasone exposure. Am J Physiol Heart Circ Physiol 322: H597–h606, 2022.

12 Organization WH. WHO recommendations on interventions to improve preterm birth outcomes. Geneva: 2015.

13 Vogel JP, Ramson J, Darmstadt GL, Qureshi ZP, Chou D, Bahl R, and Oladapo OT. Updated WHO recommendations on antenatal corticosteroids and tocolytic therapy for improving preterm birth outcomes. Lancet Glob Health 10: e1707–e1708, 2022.

14 Osterman MJK HB, Martin JA, Driscoll AK, Valenzuela CP. Births: Final data for 2020. National Vital Statistics Reports 70: 2022.

15 Magala Ssekandi A, Sserwanja Q, Olal E, Kawuki J, and Bashir Adam M. Corticosteroids Use in Pregnant Women with COVID-19: Recommendations from Available Evidence. J Multidiscip Healthc 14: 659–663, 2021.

16 Zhou CG, Packer CH, Hersh AR, and Caughey AB. Antenatal corticosteroids for pregnant women with COVID-19 infection and preterm prelabor rupture of membranes: a decision analysis. J Matern Fetal Neonatal Med 1–9, 2020.

17 Shen MJ, and Zipes DP. Role of the autonomic nervous system in modulating cardiac arrhythmias. Circ Res 114: 1004–1021, 2014.

18 Pagani M, and Lucini D. Autonomic dysregulation in essential hypertension: insight from heart rate and arterial pressure variability. Auton Neurosci 90: 76–82, 2001.

19 Ojeda NB. Prenatal programming of hypertension: role of sympathetic response to physical stress. Hypertension 61: 16–17, 2013.

20 Mizuno M, Siddique K, Baum M, and Smith SA. Prenatal programming of hypertension induces sympathetic overactivity in response to physical stress. Hypertension 61: 180–186, 2013.

21 Grippo AJ, Moffitt JA, and Johnson AK. Cardiovascular alterations and autonomic imbalance in an experimental model of depression. Am J Physiol Regul Integr Comp Physiol 282: R1333–1341, 2002.

22 Hartmann R, Schmidt FM, Sander C, and Hegerl U. Heart Rate Variability as Indicator of Clinical State in Depression. Front Psychiatry 9: 735, 2018.

23 Williams DP, Joseph N, Gerardo GM, Hill LK, Koenig J, and Thayer JF. Gender Differences in Cardiac Chronotropic Control: Implications for Heart Rate Variability Research. Appl Psychophysiol Biofeedback 47: 65–75, 2022.

24 Koenig J, and Thayer JF. Sex differences in healthy human heart rate variability: A meta-analysis. Neurosci Biobehav Rev 64: 288–310, 2016.

25 Huikuri HV, Pikkujämsä SM, Airaksinen KE, Ikäheimo MJ, Rantala AO, Kauma H, Lilja M, and Kesäniemi YA. Sex-related differences in autonomic modulation of heart rate in middle-aged subjects. Circulation 94: 122–125, 1996.

26 Carbone DL, Zuloaga DG, Hiroi R, Foradori CD, Legare ME, and Handa RJ. Prenatal dexamethasone exposure potentiates diet-induced hepatosteatosis and decreases plasma IGF-I in a sex-specific fashion. Endocrinology 153: 295–306, 2012.

27 Dixon A, and Maric C. 17beta-Estradiol attenuates diabetic kidney disease by regulating extracellular matrix and transforming growth factor-beta protein expression and signaling. AmJPhysiol Renal Physiol 293: F1678–F1690, 2007.

28 Carnevali L, Barbetti M, Statello R, Williams DP, Thayer JF, and Sgoifo A. Sex differences in heart rate and heart rate variability in rats: Implications for translational research. Front Physiol 14: 1170320, 2023.

29 Švorc P, Jr., Grešová S, and Švorc P. Heart rate variability in male rats. Physiol Rep 11: e15827, 2023.

30 Shaffer F, and Ginsberg JP. An Overview of Heart Rate Variability Metrics and Norms. Front Public Health 5: 258, 2017.

31 Shaffer F, McCraty R, and Zerr CL. A healthy heart is not a metronome: an integrative review of the heart’s anatomy and heart rate variability. Front Psychol 5: 1040, 2014.

32 Dos Reis DG, Fortaleza EA, Tavares RF, and Correa FM. Role of the autonomic nervous system and baroreflex in stress-evoked cardiovascular responses in rats. Stress 17: 362–372, 2014.

33 Team RC. R: A Language and Environment for Statistical Computing. R Foundation for Statistical Computing. 2024.

34 Pinheiro J BDRCT. Linear and Nonlinear Mixed Effects Models. R package version 3.1-164,. 2023.

35 Mastorci F, Vicentini M, Viltart O, Manghi M, Graiani G, Quaini F, Meerlo P, Nalivaiko E, Maccari S, and Sgoifo A. Long-term effects of prenatal stress: changes in adult cardiovascular regulation and sensitivity to stress. Neurosci Biobehav Rev 33: 191–203, 2009.

36 Kajantie E, and Phillips DI. The effects of sex and hormonal status on the physiological response to acute psychosocial stress. Psychoneuroendocrinology 31: 151–178, 2006.

37 Moodithaya S, and Avadhany ST. Gender differences in age-related changes in cardiac autonomic nervous function. J Aging Res 2012: 679345, 2012.

38 Kuo TB, Lai CT, Hsu FC, Tseng YJ, Li JY, Shieh KR, Tsai SC, and Yang CC. Cardiac neural regulation oscillates with the estrous cycle in freely moving female rats: the role of endogenous estrogens. Endocrinology 151: 2613–2621, 2010.

39 Tung I, Krafty RT, Delcourt ML, Melhem NM, Jennings JR, Keenan K, and Hipwell AE. Cardiac vagal control in response to acute stress during pregnancy: Associations with life stress and emotional support. Psychophysiology 58: e13808, 2021.

40 Zafar U, Rahman SU, Hamid N, and Salman H. Assessment of gender differences in autonomic nervous control of the cardiovascular system. J Pak Med Assoc 70: 1554–1558, 2020.

41 Ramesh S, Wilton SB, Holroyd-Leduc JM, Turin TC, Sola DY, and Ahmed SB. Testosterone is associated with the cardiovascular autonomic response to a stressor in healthy men. Clin Exp Hypertens 37: 184–191, 2015.

42 Hamidovic A, Van Hedger K, Choi SH, Flowers S, Wardle M, and Childs E. Quantitative meta-analysis of heart rate variability finds reduced parasympathetic cardiac tone in women compared to men during laboratory-based social stress. Neurosci Biobehav Rev 114: 194–200, 2020.

43 Nguyen P, Khurana S, Peltsch H, Grandbois J, Eibl J, Crispo J, Ansell D, and Tai TC. Prenatal glucocorticoid exposure programs adrenal PNMT expression and adult hypertension. J Endocrinol 227: 117–127, 2015.

44 Lamothe J, Khurana S, Tharmalingam S, Williamson C, Byrne CJ, Khaper N, Mercier S, and Tai TC. The Role of DNMT and HDACs in the Fetal Programming of Hypertension by Glucocorticoids. Oxid Med Cell Longev 2020: 5751768, 2020.

45 Lamothe J, Khurana S, Tharmalingam S, Williamson C, Byrne CJ, Lees SJ, Khaper N, Kumar A, and Tai TC. Oxidative Stress Mediates the Fetal Programming of Hypertension by Glucocorticoids. Antioxidants (Basel) 10: 2021.

46 Piquer B, Olmos D, Flores A, Barra R, Bahamondes G, Diaz-Araya G, and Lara HE. Exposure of the Gestating Mother to Sympathetic Stress Modifies the Cardiovascular Function of the Progeny in Male Rats. Int J Environ Res Public Health 20: 2023.

47 Jevjdovic T, Dakic T, Kopanja S, Lakic I, Vujovic P, Jasnic N, and Djordjevic J. Sex-Related Effects of Prenatal Stress on Region-Specific Expression of Monoamine Oxidase A and β Adrenergic Receptors in Rat Hearts. Arq Bras Cardiol 112: 67–75, 2019.

48 Khurana S, Grandbois J, Tharmalingam S, Murray A, Graff K, Nguyen P, and Tai TC. Fetal programming of adrenal PNMT and hypertension by glucocorticoids in WKY rats is dose and sex-dependent. PLoS One 14: e0221719, 2019.

