## Supplemental figures for "Prenatal Dexamethasone Programs Autonomic Dysregulation in Female Rats"

(A) Left Ventricle

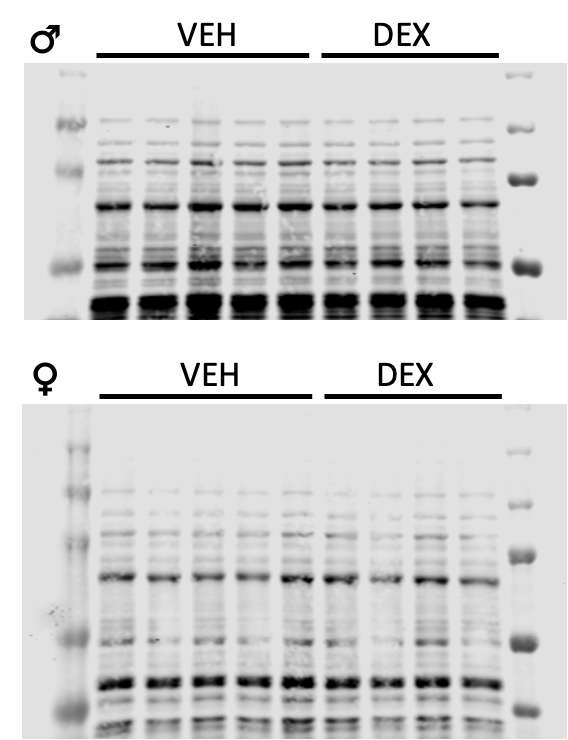

(B) Adrenal

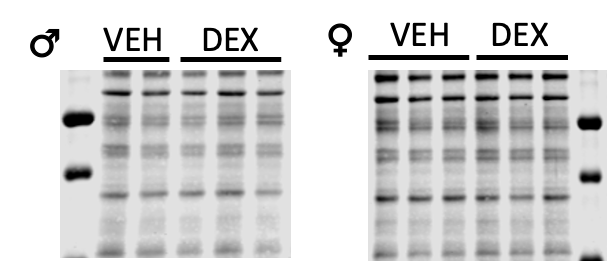

**Supplemental Figure 1.** Tyrosine Hydroxylase Western Blot Total Protein Stains for male and female left ventricle (A) and adrenal (B) blots, from Figures 7 A and B, respectively.

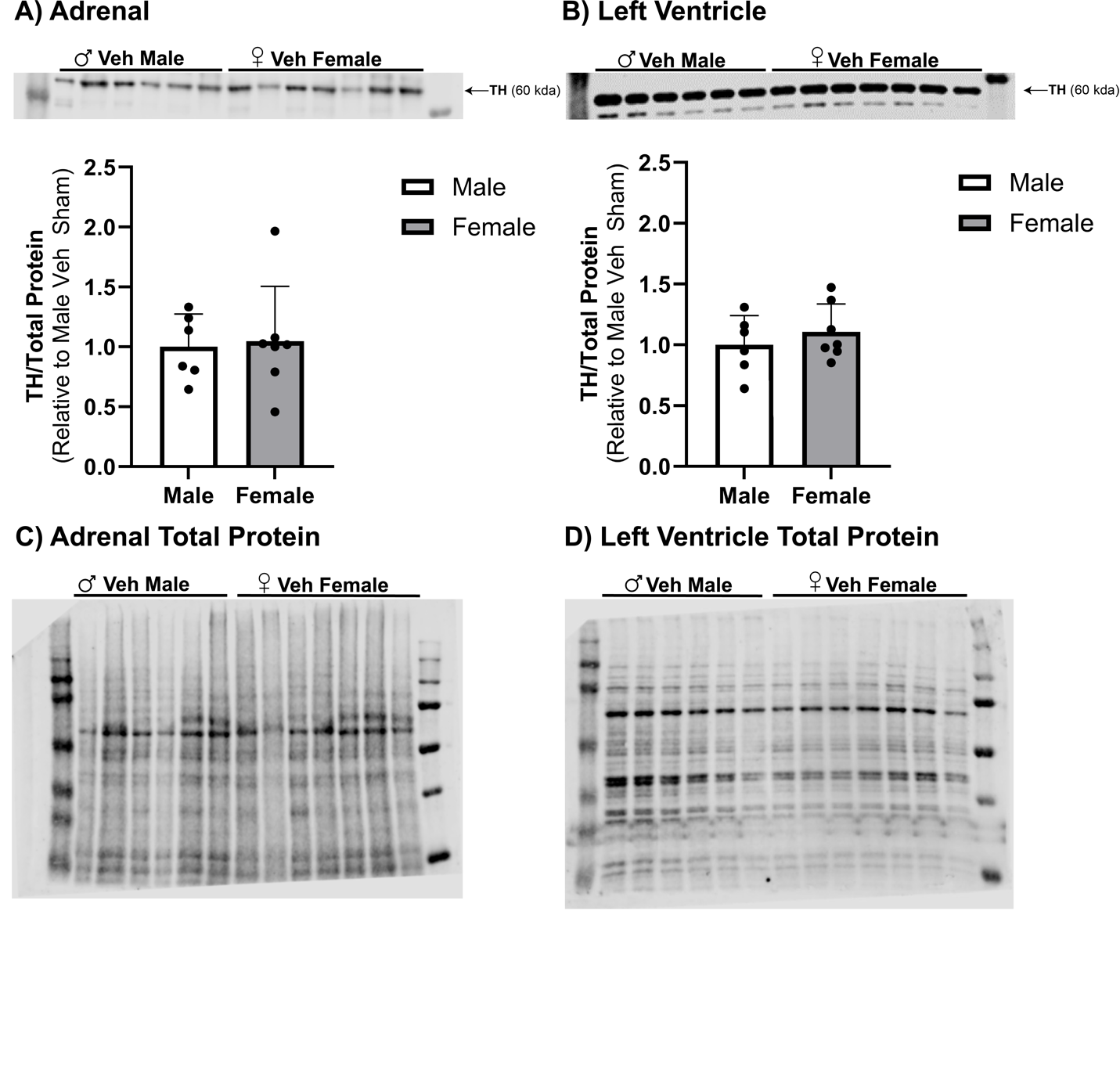

**Supplemental Figure 2.** Tyrosine Hydroxylase (TH) was assessed in adrenal (A) and left ventricle (B) from male and female offspring of vehicle treated dams. No difference in basal expression of tyrosine hydroxylase was observed between males and females for either tissue. Corresponding total protein stain for adrenal (C) and left ventricle (D).
